# COMPARISON BETWEEN SANGER, ILLUMINA AND NANOPORE SEQUENCING EVIDENCING INTRA-HOST VARIATION OF FELINE LEUKEMIA VIRUS THAT INFECTS DOMESTIC CATS

**DOI:** 10.1101/2023.11.02.563952

**Authors:** Cristobal Castillo-Aliaga, Adam Mark Blanchard, Susana Castro-Seriche, Ezequiel Hidalgo-Hermoso, Alonso Jerez-Morales, Matthew W Loose, Rachael E Tarlinton

## Abstract

Knowledge of Feline Leukemia Virus (FeLV) sequence variation has mainly been developed using Sanger sequencing methods. However, advances in next generation sequencing methods and their broad use in laboratories has been changing our understanding of viral genetics. FeLV sequencing has specific complications with the presence of both exogenous (exFeLV) and endogenous (enFeLV) virus with frequent recombination between them, limiting sequencing approaches. Here we report an FeLV-A and FeLV-B amplicon-based comparison between Sanger, Illumina, and Oxford Nanopore sequencing methods in Chilean domestic cats. We analysed the hypervariable envelope gene, where a higher number of variants as well as recombination with endogenous strains occurs. We detected multiple variants and viral quasispecies infecting the cats. We compared these three methods to evaluate the advantages and disadvantages between them. Although the Sanger method is highly reliable, it showed a high fail rate (many amplicons did not produce useable sequence) and the sequences obtained showed artificial sequence clustering when compared with the NGS methods. Illumina sequencing showed a lower error rate but could not discriminate between exogenous and endogenous viruses. Finally, Oxford Nanopore (MinION) sequencing could successfully detect low-abundance sequences and discriminate between FeLV-A and FeLV-B sequences, although its higher error-rate requires caution in interpretation of the results. Our results indicate advantages and disadvantages for each method, with the purpose of sequencing needing to be considered in the choice of method. Results of large viral phylogenetic trees combing sequences derived from mixed sequencing methods, such as those combining historical and contemporary sequencing need to be considered with some caution as artificial clustering by sequencing method may occur.

## INTRODUCTION

*Feline leukemia virus* (FeLV) is an enveloped retrovirus that belongs to the family *Retroviridae*, genus *Gammaretrovirus* (Coffin et al. 2021). The FeLV genome consists of two copies of a single positive sense RNA around 8.4 kb in length. It encompasses three genes: *gag* (capsid proteins), *pol* (reverse transcriptase enzymes, protease, and integrase), and *env* (envelope proteins). These are flanked, 5’ and 3’, by two identical un-translated regulatory sequences called LTR’s (long terminal repeats). During infection, the RNA genome is copied into a DNA genome by the reverse transcriptase enzyme and is inserted as a provirus into the host genome (Willett and Hosie 2013).

FeLV has a world-wide distribution in domestic cat populations and is one of the most important pathogens for their health (Hartmann and Hofmann-Lehmann 2020). The virus causes clinical signs such as: immune and bone marrow suppression, lymphadenopathy, lymphoma, leukaemia, oral lesions and respiratory diseases (Hartmann 2012). The virus is excreted in saliva, urine, faeces, milk and nasal secretions (Torres, Mathiason, and Hoover 2005). Social activities like grooming, sharing food and water dishes, nursing and bites during fights are the main transmission routes. It can also be transmitted via blood contact (blood transfusion or contaminated instruments) or sharing of litter trays (Little et al. 2020). Another important route of transmission is vertically, where the queen cat spreads virus to her neonatal kittens, either trans-placentally, during parturition or during nursing of kittens (Hardy et al. 1976). FeLV also has importance in wild felid health and conservation causing severe disease and mortality (Cunningham et al. 2008, Meli et al. 2010a, Chiu et al. 2019). This has been notably harmful in genetically bottle-necked populations (Chiu et al. 2019). Although, the virus can persist in wild species, domestic cats have been the infection source for non-domestic animals in the majority of cases (Meli et al. 2010b, Brown et al. 2008, Petch et al. 2022).

On the other hand, endogenous FeLV (enFeLV) are retroviruses that have integrated over millions of years into the cat genome. EnFeLV were acquired during the divergence of the Felidae and are present in domestic cat chromosomes and their close relatives in the *Felis* genus. They are transmitted from parents to descendants through chromosomes (Polani et al. 2010). Exogenous and endogenous viruses share 86% similarity at the nucleotide level with the main variation points occurring in *gag* and *env* (Chiu, Hoover, and Vandewoude 2018). The enFeLV in cat populations have multiple variations, influenced by the integration site, viral transposition events and several independent integrations during evolutionary history. Even the cats gender may influence the enFeLV quantity, due to an increased load on the Y chromosome compared to the X chromosome (Roca et al. 2005, Powers et al. 2018). Recently studies have associated higher loads of enFeLV with protective factors against exFeLV infections (Powers et al. 2018, Chiu and VandeWoude 2020). Conversely the presence of enFeLV can result in recombination events with exFeLV and produce more virulent variants (Willett and Hosie 2013).

There are six FeLV groups: FeLV-A, FeLV-B, FeLV-C, FeLV-D, FeLV-E, and FeLV-T. These arise from recombination between exogenous and endogenous viruses or mutation of exogenous viruses (Willett and Hosie 2013). The majority of the variation occurs in the *env* gene, which is the binding point between the virus and host cell (Cano-Ortiz et al. 2022). The *env* gene is composed of the Surface protein (SU) and Transmembrane protein (TM). SU includes three regions from 5’ to 3: Receptor binding domain (RBD), Proline rich region (PRR) and C-Domain. The RBD and C-Domain are involved in cellular tropism, therefore changes in these regions can modify cell receptor usage and so change clinical progression (Faix et al. 2002, Cano-Ortiz et al. 2022). More specifically, there are three main Variable Regions (VRA,VRB,VRC) identified, along with specific amino acid changes which are associated with changes in receptor usage (Boomer et al. 1997).

FeLV-A is the most widespread group that can be horizontally transmitted and often described as the least pathogenic subtype (Ahmad and Levy 2010, Biezus et al. 2023). All other groups arise de novo from FeLV-A mutation or by recombination with endogenous elements (Hartmann and Hofmann-Lehmann 2020). FeLV-B is the second most common subgroup, being detected in roughly half of all FeLV-A cases (Powers et al. 2018). FeLV-B is a recombinant between FeLV-A and enFeLV viruses and it is associated with higher numbers of neoplastic malignancies, like mediastinal lymphoma and multicentric lymphoma. Overall, FeLV-B is associated with worse clinical symptomatology and prognosis (Ahmad and Levy 2010). Although FeLV-B has historically been described as unable to be horizontally transmitted without FeLV-A, there are documented cases confirming horizontal transmission of FeLV-B without FeLV-A (Stewart et al. 2013, Chiu et al. 2019). FeLV-C, FeLV-E and FeLV-T arise from FeLV-A mutation, in general, these subtypes are associated with increased virulence but have diminished transmission ability. FeLV-C is an infrequent subgroup related to aplastic anaemia (Riedel et al. 1986), while FeLV-T is a T-cytopathic virus which has been related to acquired immune deficiency syndrome (FeLV-FAIDS) (Overbaugh 2000). FeLV-E is the last group described and was found in a natural thymic lymphoma in one cat (Miyake et al. 2016). FeLV-D is the most distant FeLV, because it is a recombinant between FeLV-A and another phylogenetically distinct endogenous retrovirus (not enFeLV), called domestic cat endogenous retrovirus (ERV-DC) (Anai et al. 2012).

Although FeLV does not have as high mutation rate as other retroviruses like Human Immunodeficiency Virus (HIV) or Feline Immunodeficiency Virus (Perelson et al. 1996, Bendinelli et al. 1995), multiple FeLV variants can be detected in a cat population or a single cat (Watanabe et al. 2013, Erbeck et al. 2021). To detect viral quasispecies variations, next-generation sequencing has been broadly used in HIV to evaluate virus variations related to drug-resistance (Teo et al. 2022). This is in contrast to FeLV studies, where the majority of *env* gene analysis have been performed using plasmid libraries and Sanger sequencing to evaluate the viral diversity (Watanabe et al. 2013, Erbeck et al. 2021, Petch et al. 2022, Biezus et al. 2023). To our knowledge, NGS in FeLV has been used in only two studies, these were: Roche 454 sequencing of an FeLV outbreak in Iberian Lynx in Spain (Geret et al. 2011) and one study using Illumina sequencing demonstrating decreased enFeLV expression in lymphomas from exFeLV negative cats (Krunic et al. 2015).

FeLV epidemiology is related to multiple factors including: household conditions, cat population management and vaccination rates, even the purchasing power parity per capita of owners can influence prevalence rates (Ludwick and Clymer 2019). Thus, developed countries generally have a controlled situation with FeLV. The current prevalence in Europe has been described as between 0.7%-5.5% (Studer et al. 2019), with a 3.1% FeLV prevalence in North-America (Burling et al. 2017). Conversely, the situation in South America, is highly variable ranging from 3% in north-eastern Brazil (Lacerda et al. 2017) up to 59.44% in Colombia (Ortega et al. 2020), whilst in Chile, the prevalence has been described as between 20.2%-33% in rural areas (Mora et al. 2015, Sacristán et al. 2021).

This study was developed in Chilean domestic cats, with the aim of evaluating the diversity of *env* gene in FeLV provirus, using the traditional sequencing method (Sanger) compared with short and long read next generation sequencing methods (Illumina and Nanopore) in order to examine sequence diversity and recombination between exogenous and endogenous FeLV with higher precision than previously possible.

## METHODS

### Samples and DNA extraction

Eight domestic cat blood samples were collected in EDTA by jugular venipuncture and kept at −20°C. All procedures and handling were done by veterinarians during clinical diagnostics in the city of Concepcion under the owner’s consent. Overarching ethical approval for this study was granted by the University of Nottingham School of Veterinary Medicine and Science Committee for Animal Research and Ethics (CARE) from the number 3672 220923.

Nucleic acid extractions were processed at the Haiken Laboratory, Concepcion, using a Geneaid® kit extraction following manufacturer’s instructions. PCR diagnosis of FeLV was made using 3’LTR primers (Cattori et al. 2006) as a first screen at Haiken Laboratory. Genomic DNA was then shipped to the School of Veterinary Medicine and Science, University of Nottingham United Kingdom.

### End-point PCR and Sample Preparation

End-point PCR using two primer pairs following methods from Erbeck *et al*, (2020) was used to amplify the envelope gene hypervariable region and 3’LTR region from FeLV-A and FeLV-B. PCR was performed using GoTaq® Long PCR Master mix (Promega) and each reaction consisted of: 12.5µl of GoTaq® Long PCR Master mix; 0.5µl (10µM) Forward primer; 0.5µl (10µM) Reverse primer; 9.5µl of Nuclease-Free water; 2µl of template DNA to reach a final volume of 25µl per reaction. The same protocol was used for each pair of primers and was performed in triplicate to confirm results. Thermo-cycler conditions used a Gene-Touch Thermal-cycler, from Bioer Technology® following the Erbeck *et al*., (2020) protocol with some modifications according to the polymerase requirements. The protocol was: initial denaturation at 94°C for 2 min followed by 45 cycles of denaturation at 95°C for 30 s; annealing at 63 °C for 30 s; extension at 72°C for 2 mins and a final extension at 72°C at 2 mins. The thermo-cycler conditions were tested with a gradient PCR to select the optimal annealing temperature and every set of PCR reaction included negative control with no template (DNA free water) and a positive control (FeLV-A DNA) which was kindly donated by Dr. Margaret Hosie, University of Glasgow. The PCR products were run by electrophoresis on a 1% agarose TAE (Fisher scientific®) gel at 80V, 400 mA for 60 minutes. DNA purification used the Nucleospin® extract II kit (Macherey-Nagel®), according to manufacturer’s instructions.

DNA quantification was performed with a Qubit 4 Fluorometer (Thermo Fisher Scientific®). FeLV-A and FeLV-B amplicons of high enough quality and quantity were selected for Illumina and Nanopore sequencing.

### Genome sequencing

Amplicons from 8 domestic cats were sequenced by three different methods detailed in Table 1 and all results sequencing were submitted to GenBank. Five amplicons were sequenced by Sanger method in a commercial laboratory (Eurofins Inc., Germany).

**Table 1.**
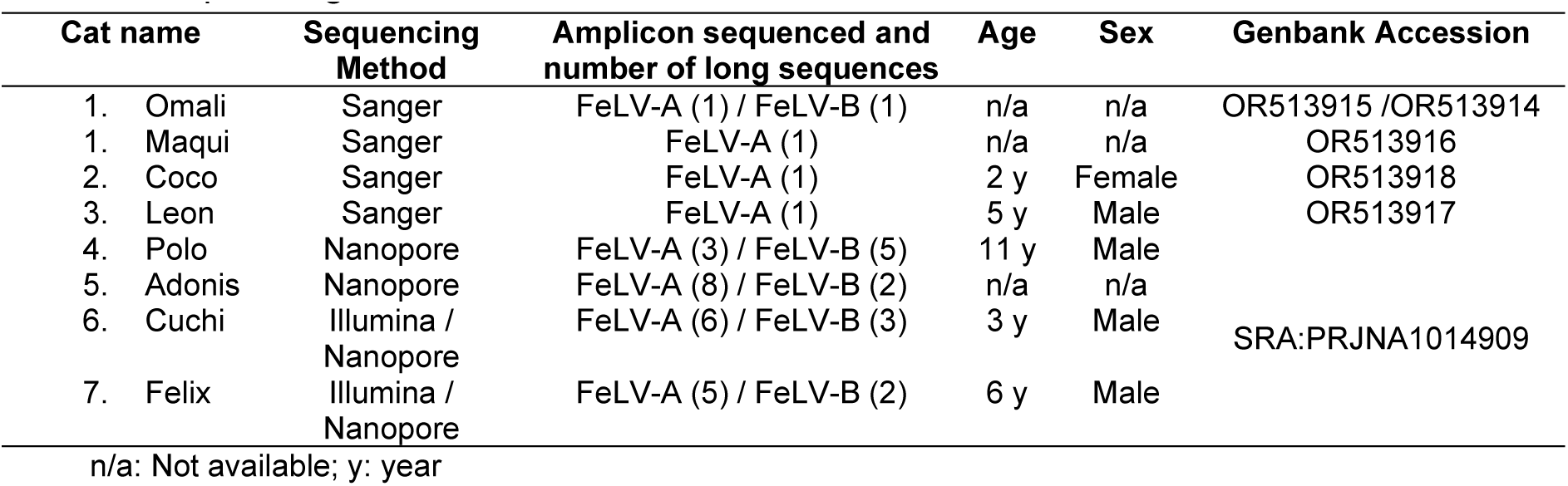
Sequencing method in Chilean domestic cats.

For Illumina sequencing, amplicons of FeLV-A and FeLV-B were pooled and sequenced using the Illumina platform (HiSeq4000) with 300-bp paired-end distances using a coverage of 30x by Novogene Europe, Cambridge, United Kingdom.

The libraries for Nanopore sequencing were processed following the Native Barcoding Kit 24 V14 protocol (Oxford Nanopore Technologies) with FeLV-A and FeLV-B amplicons individually barcoded. Library Quantification was carried out using a Qubit fluorometer and the Qubit dsDNA HS Assay kit (Thermofisher). 20 fmol of DNA library was loaded onto a R10.4.1 flow cell and sequenced using an Oxford Nanopore MinION Mk1C.

### Read Processing, Assembly and Mapping

Illumina reads were trimmed and adaptors removed using FastP v0.23.1 (Chen et al. 2018). The Nanopore raw reads were base called using Guppy v6.4 and super accurate model was used. and Nanopore adaptors were eliminated using Porechop v0.2.4 (Wick et al. 2017), NanoFilt v2.6.0 (De Coster et al. 2018) was used to filter and discard reads shorter than 500 and longer than 4,000 bp with a quality of <Q10.

De Novo Assembly from Illumina reads was performed using MEGAHIT (Li et al. 2015) and for Nanopore reads, Flye (Lin et al. 2016) was used for FeLV-A barcodes. FeLV-B barcodes did not assemble reliably in Flye due to a mix of endogenous and exogenous sequences.

The contigs generated by MEGAHIT and raw reads from Nanopore, were mapped to FeLV-A FAIDS (M18247), enFeLV (M25425, LC196053), FeLV-T (M89997); FeLV-C (M14331), (MT302119, MT302153 and MT681671), Gardner-Arnstein FeLV-B (V01172) and FeLV-A and FeLV-B Chilean Sanger sequences. Illumina sequences were mapped using Bowtie2 (Langmead and Salzberg 2012). Nanopore reads were mapped using Minimap2 (Li 2018) with two preset options: the first was all raw reads by the more sensitive preset options “Oxford nanopore reads” (map-ont). The second approach was mapping to specific variations of interest (Chilean Sanger, FeLV-T, FeLV-E and US FeLV-B) with raw reads and using the preset options for “long assembly increased the specificity up to 5% sequence divergence” (asm5) to consider those under-represented sequences that could be missed by Flye. Reads were considered in the second approach if there were at least 30 reads with similar variations to generate a consensus sequence.

All contigs and consensus sequence were extracted and imported to Geneious v2023.0.4 (Biomatters Inc., Newark, NJ) to annotate and continue phylogenetic analysis.

### Phylogenetic Analysis of FeLV

The dataset was constructed in Geneious (Biomatters Inc., Newark, NJ) using all available complete *env* gene FeLV related sequences and were annotated and aligned with Muscle (Edgar 2004). This was exported and used to construct phylogenetic trees and evaluate recombination between FeLV-A and enFeLV sequences.

The alignment was sent to the IQtree web server (Trifinopoulos et al. 2016), where a maximum likelihood tree was inferred using the ModelFinder (Kalyaanamoorthy et al. 2017) to select the best-fitted model nucleotide substitution. The model chosen was general time reversible (GTR+F+G4), branch support analysis was ultrafast with 5,000 bootstrap (Hoang et al. 2018). EnFeLV was used as an outgroup to root the tree and the phylogenetic tree was visualized in FigTree v1.4.4 (Rambaut 2018).

### Intra-host Variation Analysis and Recombination Analysis

The intra-host single nucleotide variation (iSNV) was done using BWA-MEM (Li 2013) and iVar (Grubaugh et al. 2019). All libraries were mapped to FeLV-A reference sequence (M18247) using BWA-MEM and a consensus sequences were generated using iVar consensus command for each library. All reference sequences generated by iVar were cut at nucleotide 66 (M18247), to initiate at the same nucleotide position. The output was used to call single nucleotide variants and indels in iVar. The minimum quality parameter was <Q10 and the minimum frequency selected was 0.01 and was considered as true only those statistically significative (p <0.05).

Recombination analysis was carried out in RDP v4.101 (Martin et al. 2015) using six different methods and default parameters. The methods used were RDP (Martin et al. 2015), GENECONV (Padidam, Sawyer, and Fauquet 1999), Chimaera (Posada and Crandall 2001), MaxChi (Smith 1992), Bootscan (Martin and Rybicki 2000), 3Seq (Boni, Posada, and Feldman 2007) and Siscan (Gibbs, Armstrong, and Gibbs 2000). Recombinants were considered if at least three methods detected breakpoints. The analysis included the reference sequences used previously and all sequences obtained by the three methods.

## RESULTS

Amplicons for FeLV-A PCR showed lengths around the expected amplicon size (∼1.9kb) with some products showing multiple bands. All FeLV-B PCR amplicons had similar lengths to the expected amplicon (∼1.8 kb). Additionally, some FeLV-B PCR products showed the presence of an extra band below 1.8 kb (these were identified as truncated enFeLV in the NGS results).

Twenty Sanger sequencing (10 of FeLV-A and 10 of FeLV-B) of isolates were performed to confirm PCR product identity. Only four samples of FeLV-A were successfully Sanger sequenced (Omali, Maqui, Coco and Leon) with around 1.8 kb of sequence obtained, these DNA samples however did not have enough volume remaining to sequence with the other methods. BLASTn analysis from NCBI showed 97-99% identity with FeLV-FAIDS (M18247.1) sequence. Only one FeLV-B sample (Omali) was sequenced successfully (1,793bp) by the Sanger method, showing 98% identity with FeLV-B sequences from the USA (MT302107.1). Most amplicons failed to produce good quality results with the Sanger method, due to high interference from the presence of multiple variants.

For the amplicons sequenced by Illumina, one long sequence per cat was assembled with multiple small contigs in different gene sections. Four long sequences between FeLV-A and FeLV-B were obtained and were considered in the alignment for further phylogenetic analysis. For Nanopore sequencing, two contigs per cat were obtained by the first approach (De Novo assembly using Flye). These contigs were around 1,900bp of length. Afterwards, using the second preset options in Minimap2, 13 consensus sequences were obtained (6 sequences from Adonis; 4 from Cuchi; 5 from Felix; 1 from Polo). These were between 1,764bp to 1,602bp of length, including insertions, deletions and missed nucleotides in the sequence start.

The alignment of all these FeLV-A included 29 sequences belonging to 8 different cats. The alignment shows a similarity between 100-94.8% at nucleotide level. Amino acid changes were mainly identified in the Surface unit (Figure 1 and supplementary information). Seven variations were present in almost all Chilean sequences, 5 variations showed 2 possible amino acids (Supplementary table). Other variations were less often detected, these were mainly present in Sanger sequences. The most distant sequence was a Sanger sequence (Omali_S). Illumina amino acid changes were consistent with Nanopore results, except for some minor changes in Felix_5_I in positions 202 and 230 (Supplementary information). The main amino acid insertions were detected in Maqui (Sanger). Deletions were observed in the SU region from nanopore sequences generated by consensus in Felix (5_Cs_Np), Cuchi (4_Cs_Np) and Adonis (7_Cs_Np).

**Figure 1.**
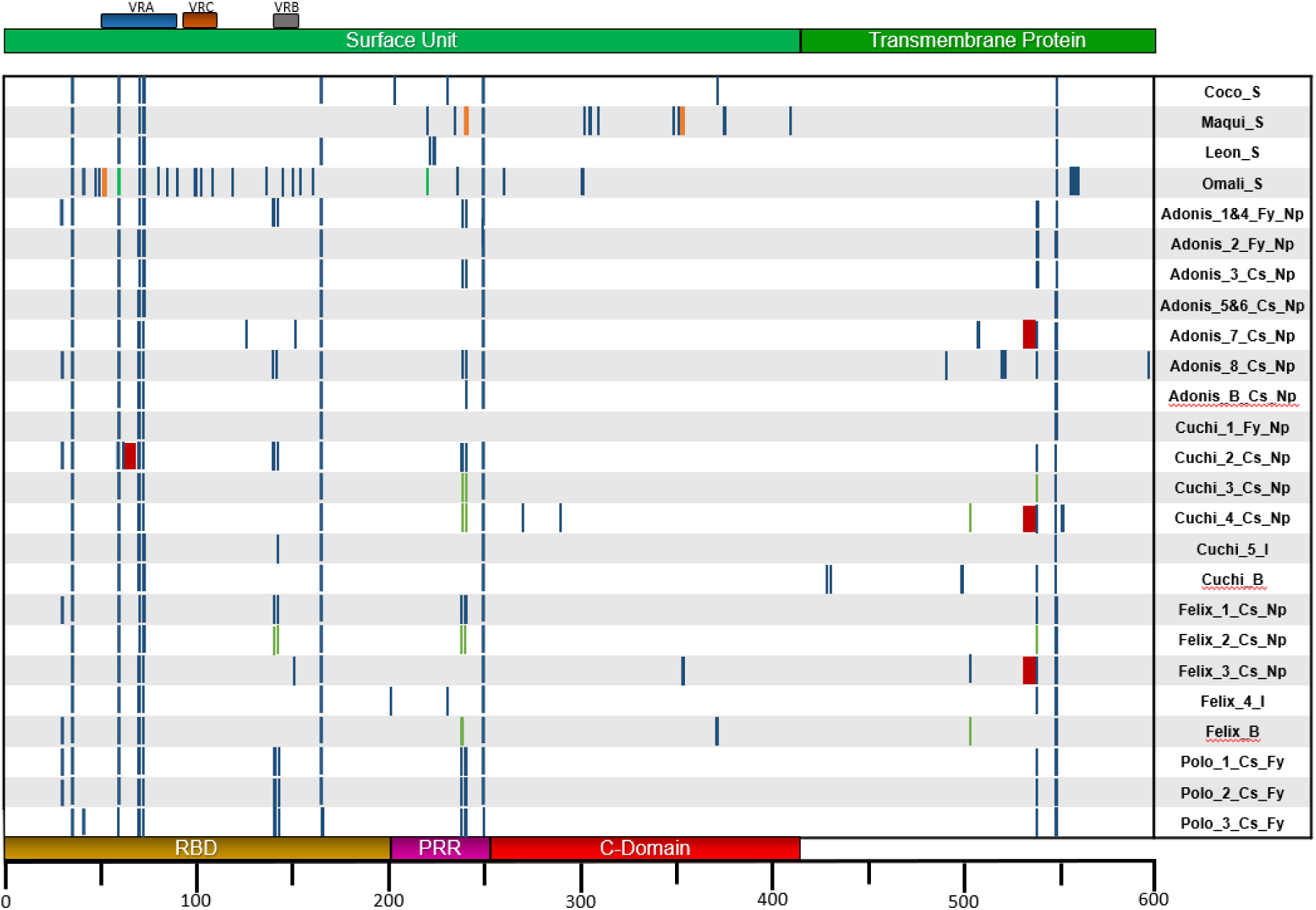
Scheme of alignment of envelope gene showing amino acid changes positions in FeLV-A sequences obtained from Sanger (S), Illumina (I), and Oxford Nanopore (Np) method. The coloured horizontal bars show the variable regions (VRA, VRC and VRB), the surface unit (light green), Transmembrane protein (green), Receptor binding domain (RBD) (yellow), Proline rich region (purple) and the C-Domain (red). Each blue vertical bar shows a single amino acid variation compared to the reference sequence, broader bar indicates two or more adjacent variation. Red vertical bars are deletions; Orange bars are insertion; Green bars alternate amino acid in the same position. Sequences are labelled by cat name, sequencing methods, assembly method, consensus (Cs) or De novo assembly in Flye (Fy).

The FeLV-B amplicon in Illumina only produced one long sequence and multiple small fragments concordant with FeLV-B or enFeLV. The longest sequence was obtained from Cuchi.

From Nanopore FeLV-B amplicons, 17 sequences were generated, although 4 of them were considered as FeLV-A. The other sequences were closer related to enFeLV. The longest sequence was 1,764bp and the shortest was 1,474bp. These short sequences were missing nucleotides at the beginning of the sequence (5’ end). The percentage similarity was 99.5% to 82.3%. The Illumina sequence was the most distant phylogenetically and some products were highly homologue with FeLV-A.

The recombination analysis performed in RDP4 showed three recombination breakpoint sites which were unique in the FeLV-B amplicons. All recombination breakpoints were derived from enFeLV genes with variable length inserted in the SU. The longest enFeLV insertion was in Cuchi_FeLV-B_I encompassed the whole RBD and partial PRR. While the second pattern was a smaller portion of enFeLV inserted in the end of RBD (Figure 2).

**Figure 2.**
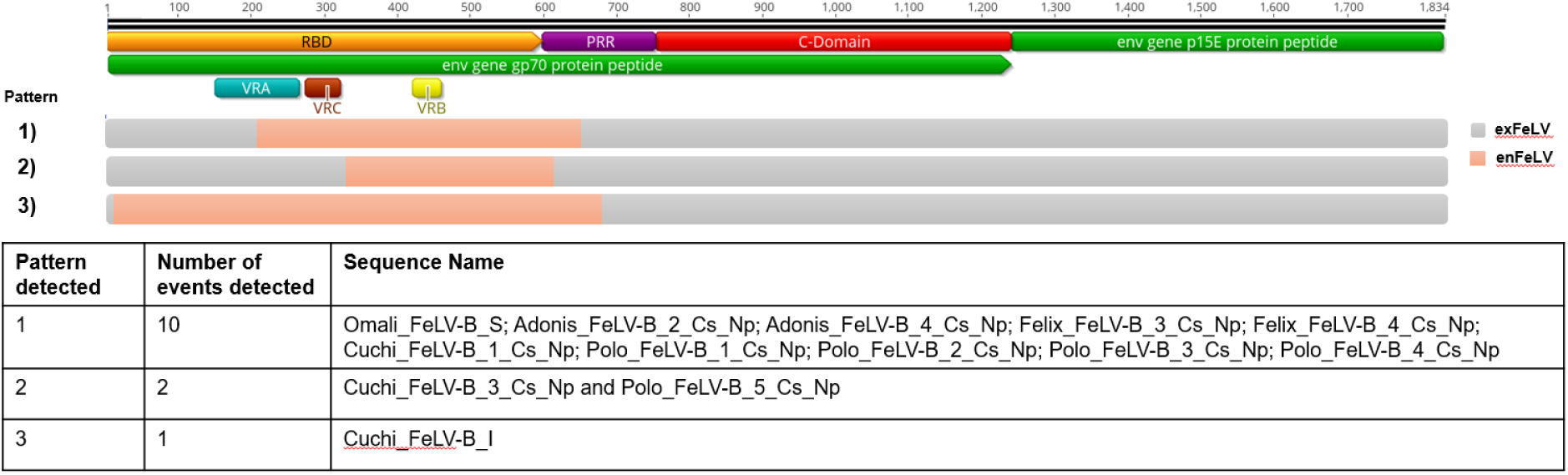
Scheme of recombinant patterns from RDP4 analysis. The reference sequence is the envelope gene from FeLV-A_FAIDS (M18247) sequence annotated in Geneious 2023.1.1. All sequences were considered in the analysis, but recombination was only detected in FeLV-B amplicons. Sequences from Sanger (S), Illumina (I), and Nanopore (Np) are shown. The gray bar color is sequencesrelated to exogenous FeLV (exFeLV) and orange is for endogenous FeLV (enFeLV). VRA, VRC and VRB are the variable region A, B and C. RBD is the receptor binding domain, PRR: Proline rich region. The table enumerates the pattern detected, and number of sequence where were detected the recombination and sequence name when were detected.

A total of 174 sequences were considered in the phylogenetical analysis, including all *env* complete sequences from GenBank (in supplementary information) and a second tree was constructed including 104 sequences (Figure 3), including phylogenetically closer sequences and representative clades. The tree is rooted in enFeLV, being subdivided into two major clades, enFeLV and FeLV-A. The Chilean FeLV-A clade is clearly identified next to the US (United States of America) sequences and the cats are homogenously represented through the Chilean clade. Sanger sequences are more related to each other than the rest of the NGS sequences. In contrast to the FeLV-A Chilean clade, the FeLV-B sequences are not closely related in a unique clade, where the biggest nanopore group are located next to sanger sequences and the Illumina sequence is closer to Gardner-Arnstein (V01172) with a third clade formed by Cuchi_FeLV-B_3_Cs_Np and Polo_FeLV-B_5_Cs_Np, closer to FeLV-A. This third clade also demonstrated the second recombination breakpoint pattern with a shorter enFeLV portion.

**Figure 3.**
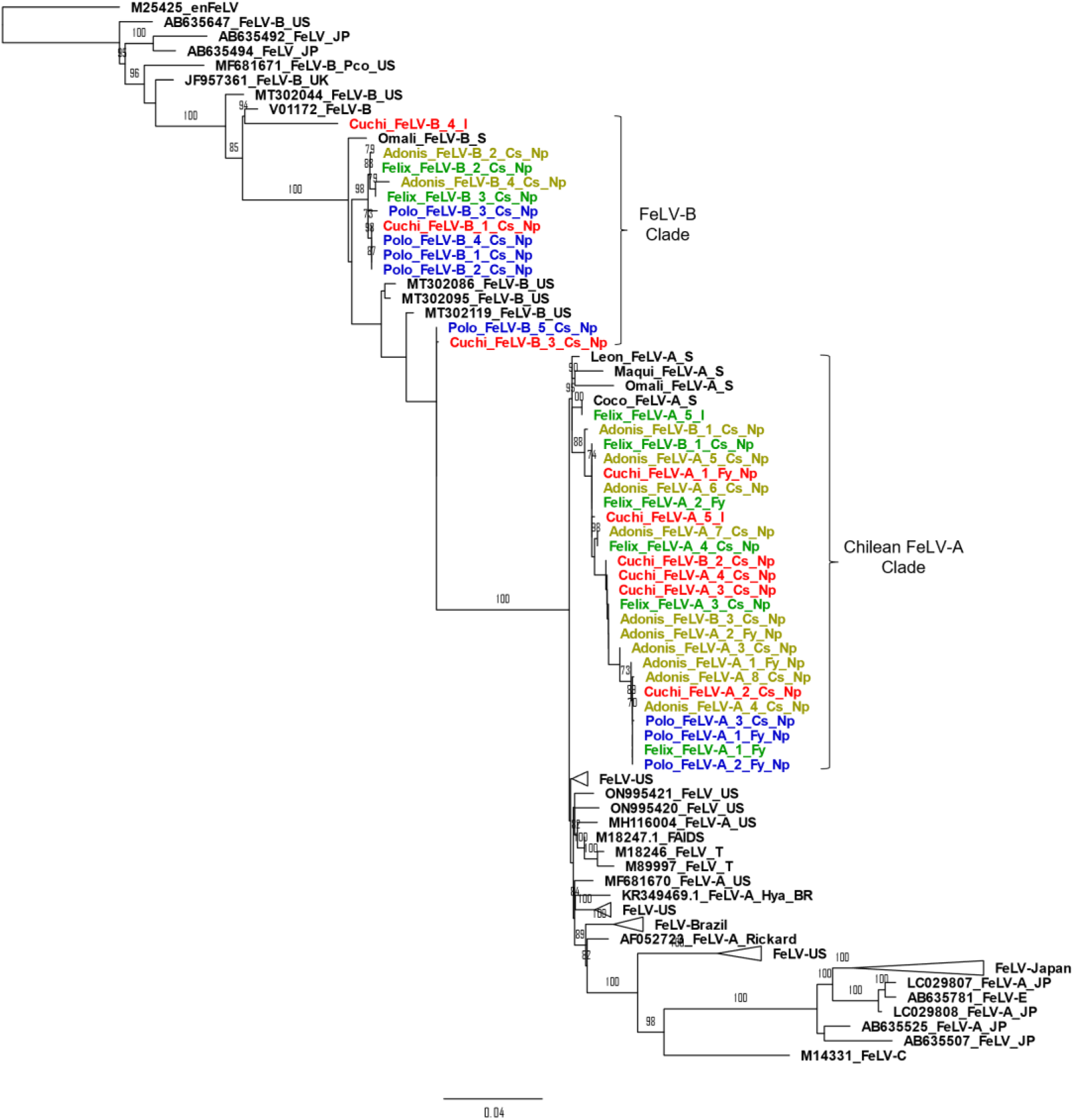
Phylogenetic tree of nucleotide env gene constructed with 5000 bootstrap approximation and rooted in enFeLV (M25425). Eighty-four FeLV sequences were considered, including references sequences of FeLV-A, FeLV-B, FeLV-C, FeLV-T and FeLV-E. Clades from the US and Japan have been collapsed for clarity (horizontal triangles). The sequence name colour indicates the cat sampled (Cuchi in red; Polo in blue; Felix in green; and Adonis in yellow). Sequences obtained from GenBank were called according to its reference number, type of FeLV (if is known it), species (if was required), and country of origin. The sequences obtained in this study were called according to name; FeLV amplicon obtained; number sequence per cat; sequence obtained by consensus (Cs) or de novo assembly in Flye (Fy); and sequencing method, Illumina (I), Nanopore (Np) or Sanger (S).

iVar intra-host variation analysis was applied to all NGS results, comparing the two different NGS methods (Cuchi and Felix) and comparing FeLV-A and B amplicons from nanopore results (Figure 4). Differences between the methods and amplicons were observed (Complete tabulate tables are in supplementary information). In general, the nanopore results in FeLV-A amplicons detected around 400 variation events, but only a small portion of these were statistically significant (less than hundred). Similarly, in FeLV-B above of 800 variations events were detected, but less than 200 on average were statistically significant. This contrasted with the Illumina results, where less than 100 variations were observed and half of them were considered as true. All Illumina variations were detected in low frequencies and were located from the end of RBD and beginning of PRR. Nanopore results showed a longer distribution of variation through all the *env* gene from FeLV-A amplicons. The cat “Felix” FeLV-A amplicon showed high frequencies of variations in the RBD, but these were highly similar to the FeLV-B amplicon from the same cat. All FeLV-B amplicons showed a high number of variations in the first portion of RBD and a minor group of changes in TM (Figure 4).

**Figure 4.**
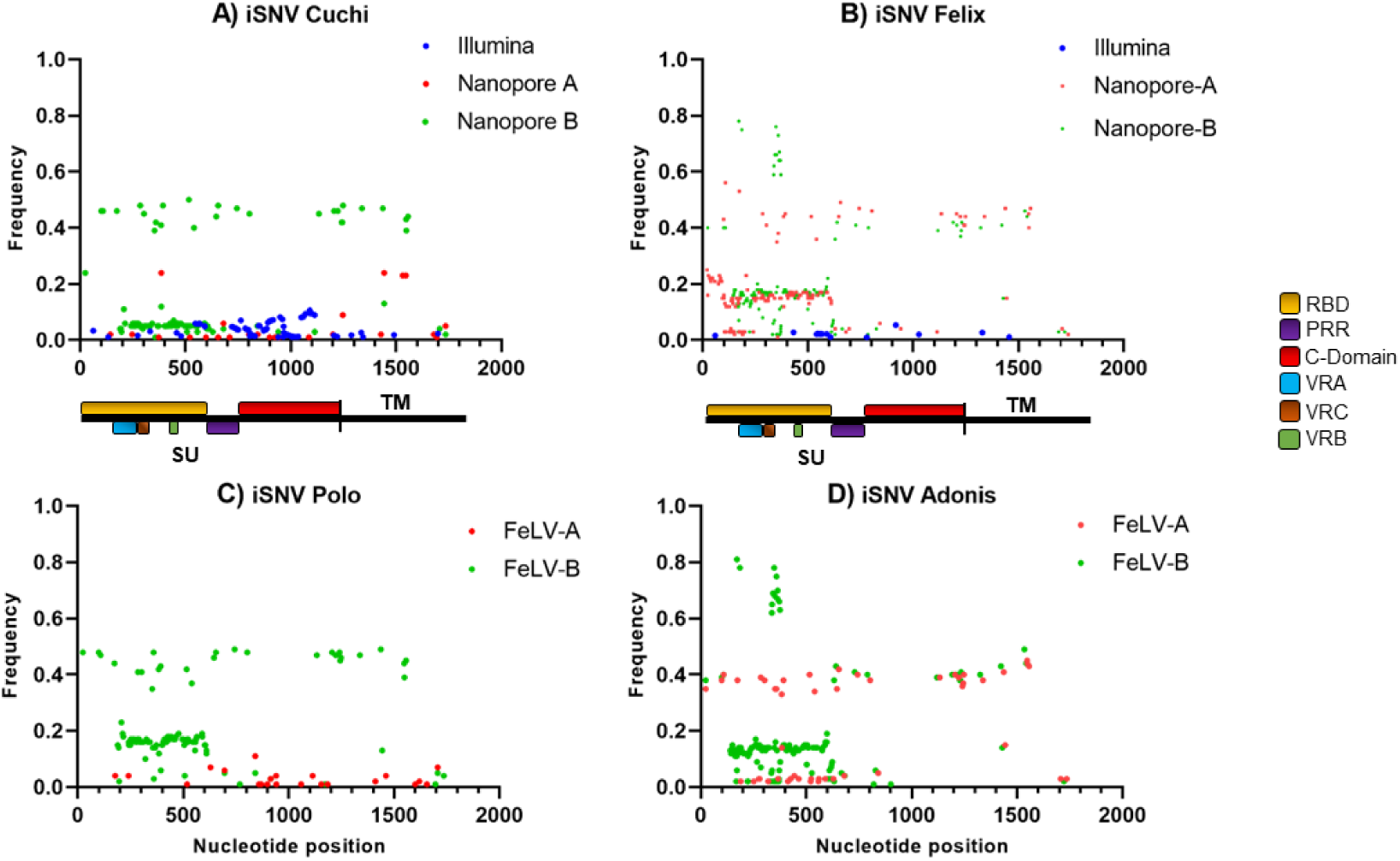
iSNV sites detected in env gene amplicons from NGS results. iVar software was used to generate a consensus sequences per library and to be used as reference sequence (∼1,800bp). In the middle are schemes of FeLV-A (M18247) showing gene positions and main variation sites (RBD: Receptor Binding Domain; PRR: Proline Rich Region; VRA, VRC and VRB are variable region A, B, and C respectively. Each graph (A,B, C and D) represents the cat sampled and are compared frequency and nucleotide position classified by amplicon and sequencing method. The blue points correspond to variation points by the Illumina method (in A and B graph), the red points are variation points in FeLV-A amplicon and green points are FeLV-B amplicon variations (all graphs).

With regards to nucleotide variation (Figure 5) compared to the reference sequence (1765bp), its frequency is led by adenine (29.35%) and cytosine (27.20%), followed by thymine (21.53%) and guanine (21.93%). The most common variation in Cuchi and Polo (all libraries) were A→G and T→C. Similarly, Felix-I has higher variation to A→G and T→C, although nanopore results had a similar variation percentage between A→G and T→C with C→T and G→A. Adonis, in both nanopore libraries, showed an increased percentage of C→T and G→A mutations.

**Figure 5.**
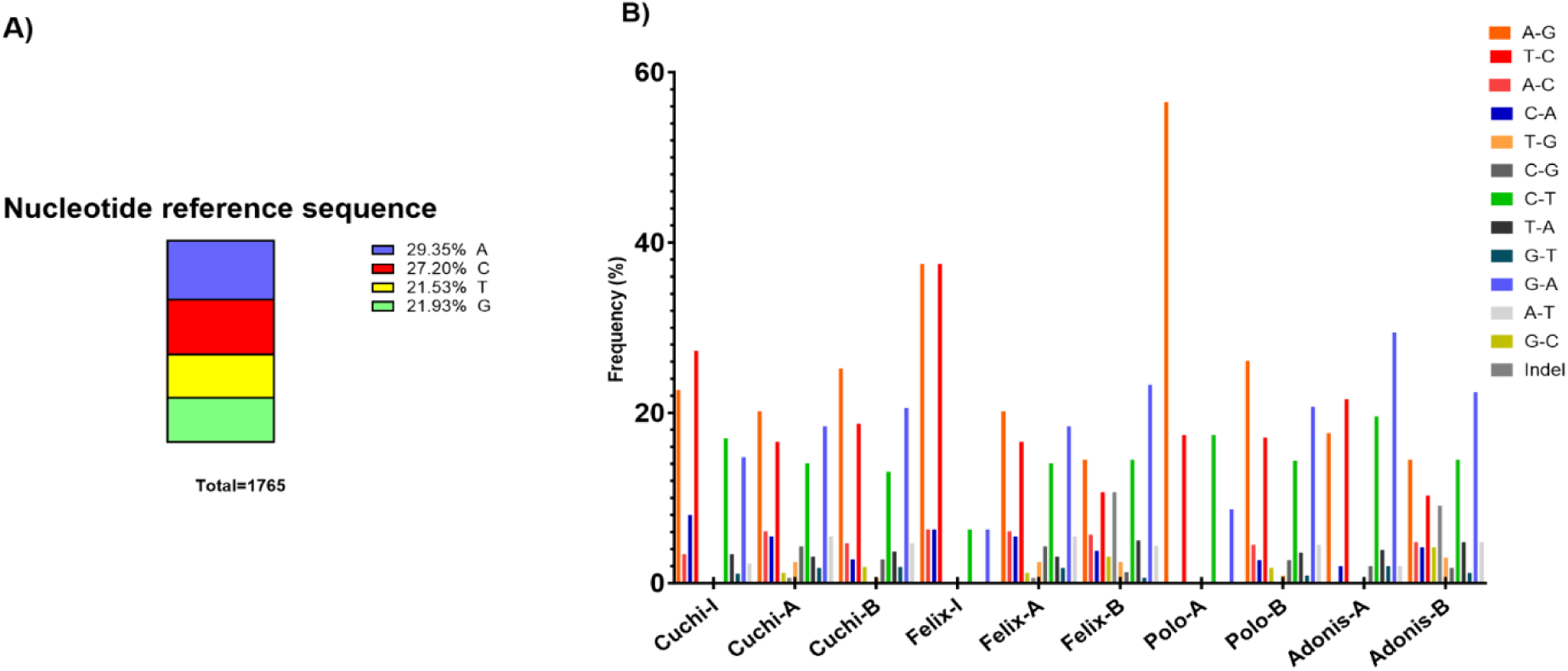
Nucleotide variation in sequences obtained from NGS sequencing. A) Graph of nucleotide frequencies in reference sequence. B) Bar graph showing nucleotide change frequency. Each colour represents one nucleotide change and there are grouped by cat and amplicon obtained. Nanopore results are pointed out the cat and from which amplicon was obtained, FeLV-A (A) or FeLV-B (B). Illumina results are represented with the cat name and “I”.

## DISCUSSION

Originally, FeLV variant characterization has been studied by viral interference assays in cell culture but genetic advances have facilitated the evaluation of specific genes and correlates with functional proteins (Chiu, Hoover, and Vandewoude 2018). The most common genetic analysis in FeLV has been based in the LTR due to its high conservation. Nonetheless, the importance of viral recombination has driven analysis of the envelope gene. Studies using molecular analysis based in *env* are available only in a few countries: Japan, the US (Erbeck et al. 2021), the UK (Stewart et al. 2013), Spain (Geret et al. 2011) and Brazil (Cano-Ortiz et al. 2022, Biezus et al. 2023). This is mainly because the work is laborious, with multiple steps such as end-point PCR or nested PCR, plasmid cloning and Sanger sequencing (Watanabe et al. 2013, Erbeck et al. 2021). Although NGS has had a rapid advancement, these techniques are still new in FeLV diagnosis, where the main problem is the high homology between enFeLV and exogenous FeLV (Chiu 2018). All our sequencing results were broadly consistent, however there were differences detected depending on the sequencing method. Here we documented the comparison of *env* gene amplicon-based sequencing using three methods and evidencing advantages and limitations of each technique.

Sanger sequencing has been the most widely used method in *env* analysis, it is highly reliable but only can generate only one single read per amplicon. To obtain clear reads in a mixed population (like a retroviral quasispecies), it is usually necessary to create plasmid clones and sequence these individually (Watanabe et al. 2013, Erbeck et al. 2021). Our results were performed directly from the PCR amplicon because it was not the aim of this study and, as expected, most sequences did not show good quality. Only those with clear high-quality reads were considered in this methods comparison. Similarly, FeLV-B amplicons had worse quality due to the presence of FeLV-A and enFeLV in the same amplicon (as was confirmed by the NGS results). The main advantage of the Sanger method is the lower error rate, described as 0.001% (Hoff 2009), however added to this error rate should be considering the multiple steps, PCR amplifications, plasmid cloning and sequencing (Cheng, Fei, and Xiao 2023). Besides, studies based on HIV has shown that Sanger sequencing has a limited capacity to detect low-abundance intra-host variants when these are under a threshold of 20% of frequencies. This could mean missing information about less-representative but still clinically important sequences (Ávila-Ríos et al. 2020). Our FeLV-A Sanger sequences clustered closer together than sequences from the other two methods with only one sequence per cat. These sequences, however, had a higher number of SNPs and indels per sequence than contigs obtained by other methods, with the “Omali” Sanger sequence showing the highest number of amino acid variations. This likely reflects the consensus nature of the Sanger sequences, but also there was a high rate of failure for Sanger sequencing (only 5 of 20 sequences attempted showed good quality), likely reflecting the sequence diversity of the PCR amplicons, failing to produce a clean enough sequence to read.

Illumina short-read method is described as the most reliable method for detecting iSNVs, showing only slightly higher base-pair error than Sanger sequencing, between 0.26%-0.8% (van Dijk et al. 2014). However, it was not possible to assemble many long contigs from the Illumina short read data. In order to increase the number of contigs assembled, various bioinformatics modifications were tried, for instance reducing the filtering options in FastP and using SPAdes as an assembler, but this did not make a substantial difference to contig recovery. Only one Illumina FeLV-B sequence was therefore available for recombinant analysis with this sequence a different structure to those detected by the Sanger or nanopore methods. Due to a failure to generate long contigs, it was only possible evaluate SNVs using the iVar software comparing reads mapped to reference sequences of FeLV. Using iVar software a reduction in the minimum frequency threshold to include under-represented reads was required, because a high number of artifacts endogenous FeLV from the cat genome were detected as well.

Oxford Nanopore was a successful tool to discriminate between FeLV-B and FeLV-A and to determine the structure of the enFeLV and exFeLV recombinants in the FeLV-B PCRs, but it is a still developing tool and its error rate is described as between 5-15% (Rang, Kloosterman, and de Ridder 2018). Although the successive MinION versions have been decreasing this error rate. At the iSNVs level, its results should be cautiously treated in order to minimize the error (Grubaugh et al. 2019). This is reflected in the sequences retrieved in this study by the number of sequences with X (uncertain amino acid) reads. The Nanopore long read method is however much better suited to detecting large indels in sequences and this is reflected in this study by its better success at retrieving and mapping FeLV-B recombinants. Similarly to the main FeLV-B recombination pattern previously described (Cano-Ortiz et al. 2022), the majority of sequences obtained included RBD and portion of PRR derived from enFeLV. Nonetheless, we used a conservative criterion to generate consensus sequences. Some were in fact exogenous reads with a longer portion derived from enFeLV, where the exogenous sequence had an endogenous insertion encompassing the RBD up to almost the whole C-Domain, but these were detected in a small number of reads therefore were not considered in further analysis.

Regarding amino acid and nucleotide variations, some of them were present in all Chilean reads when compared to the reference sequence (a US isolate from 1988) and are clearly features of Chilean isolates. Other iSNVs were present in only a small number of sequences at a low frequency and probably represent minor variants within the quasispecies mix. Interestingly many of these iSNVs were outside what are usually regarded as the more variable regions of the *env* gene. Different recombination patterns are also evident in the sequences. A comprehensive FeLV recombination review (Cano-Ortiz et al. 2022) showed multiple recombination patterns between exogenous and endogenous FeLV. In our data recombination’s between enFeLV and exFeLV were frequent in the RBD with very few in the C-Domain. This is reflected in the lower number of SNPs in the C-Domain with most related to exFeLV variations (as few FeLV-B isolates contain enFeLV sequence in this domain. Our results confirm the previous data with the RBD (the most important region for binding to the receptor) the most variable site and the site of most frequent recombination with enFeLV. The C-Domain appears to be the most highly conserved region of the virus. This domain is known to play a role during viral infection as a secondary receptor, particularly for the FeLV-C strain, and appears to be particularly important in tropism for particular receptors/cell types (Boomer et al. 1997, Ramsey, Spibey, and Jarrett 1998, Gwynn et al. 2000, Rey, Prasad, and Tailor 2008). The lack of sequence variation in this study indicates that conservation in this domain is clearly critical to the virus.

The NGS results also showed that neither the PCR primers nor the mapping software completely discriminated between FeLV-A and FeLV-B sequences. FeLV-A sequences were generated from FeLV-B amplicons, and during the iSNV analysis, all sequences were mapped to a consensus generated from FeLV-A reads, however, in Adonis-A and Felix-A, these showed a group of high frequency iSNVs closely related to FeLV-B. The pattern of nucleotide change (A→G, T→C, and A→T) is similar to that in a study of FeLV cats infected with tumors (Rohn et al. 1994). This type of mutation signature is commonly caused by the innate anti-viral APOBEC protein system, one of the main host responses in retrovirus infection (Stavrou and Ross 2015). These proteins generate hypermutated viral genomes with an increased number of G→A mutations to produce non-infective virions (Terry et al. 2017) and our results are consistent with this.

Each sequencing method produced different results even for the same animal and this needs to be considered when comparing large phylogenetic trees generated from amplicons derived from different sequencing methods. Each sequencing method has advantages and disadvantages, but phylogenies based on Sangar sequencing probably overestimate viral variation and may result in artificial sequence clustering when compared with the NGS methods. Moreover, other factors, like the input DNA amounts and concentrations used need to be considered in interpreting viral sequence variation. Our Illumina analysis was done using the minimum DNA requirement (around 5µg) and using 30x of coverage input, conversely, we used the maximum allowed DNA input for nanopore (around 250µg), this may have influenced the sequencing quality and under-represented sequences in Illumina as has been reported before (Illingworth et al. 2017) contrasting to the higher diversity detected in nanopore, although similarly to other reports, minority reads were masked and diluted by a greater number of sequences during consensus read construction (Mori et al. 2022).

Overall, the different methods were useful for different purposes with Sanger and Nanopore less reliable for iSNVs detection than Illumina, on the other hand, Nanopore was more successful for recombinant FeLV-B detection. With the growth in sequencing studies using different technologies, it will become increasingly important to consider which sequencing method was used in interpreting geographic and temporal variations in viral sequences, as well as designing and standardizing appropriate pipeline analysis to improve the quality of sequence, data interpretation and avoid sequence dilution, effects as has begun to be implemented in diagnostic HIV sequencing (Ávila-Ríos et al. 2020, Lee et al. 2020, Mori et al. 2022). Artificial clustering and estimation of sequencing variation due to sequencing artifact may complicate the interpretation of large viral phylogenies, particularly in viruses like FeLV where there is high virus circulation and recombination. Due to this being the first study conducted with NGS sequencing strategies for, there are no previous comparable data. Future works should include a greater number of animals and geographically distant FeLV, to analyse the viral diversity in domestic and non-domestic felids.

## ACKNOWLEDGEMENTS

The first author gratefully acknowledges ANID (Agencia Nacional de Investigacion y Desarrollo, Chile) – Scholarship ID 72210211 for the support to this research.

