## Supplementary 1 for "COMPARISON BETWEEN SANGER, ILLUMINA AND NANOPORE SEQUENCING EVIDENCING INTRA-HOST VARIATION OF FELINE LEUKEMIA VIRUS THAT INFECTS DOMESTIC CATS"

### SUPPLEMENTARY INFORMATION

**Table SP. 1.** Alignment of env gene showing amino acid changes in FeLV-A sequences obtained from Sanger (S), Illumina (I), and Oxford Nanopore (only shows number) methods. The variable regions VRA, VRB, VRC, proline rich region (PRR) and C-domain are shaded. Variations from the reference sequence (FeLV-FAIDS Genbank number M18247) are shown in red. Sequences are labelled by cat name and sequence number. Uncertain amino acids are pointed out with a X.

```
             |20                           |50                                                |100                                              |150
FAIDS        QTNTQANATSMLGTLTDVYPTLHVDLCDLVGDT-WEPIVLSPTNVKHGARYPSSKYGCKTTDRKKQQQTYPFYVCPGHAPSLGPKGTHCGGAQDGFCAAWGCETTGEAWWKPSSSWDYITVKRGSSQDNNCEGKCNPLILQFTQKGKQAS
Coco_S                       .A...............-......N..........SD.............................................................................................R...
Leon_S                       .A...............-......N..........SD.............................................................................................R...
Maqui_S                      .A...............-......N..........SD.................................................................................................
Omali_S                      .A....Y......A.E.PG.....D..........SD.........K...P......L.......V.S...L..........L...................L.......R........H.......H......
Adonis_1&4      .......I......A...............-......N..........SD....................................................................R.E......................R...
Adonis_2        ..............A...............-......N..........SD.............................................................................................R...
Adonis_3        ..............A...............-......N..........SD.............................................................................................R...
Adonis_5&6                         ...........-......N..........SD.............................................................................................R...
Adonis_7                           ...........-......N..........SD.......................................................X.......................X.............R...
Adonis_8        .......I......A...............-......N..........SD....................................................................R.E......................R...
Adonis_B        ..............A...............-......N..........SD.............................................................................................R...
Cuchi_1         ..............A...............-......N..........SD.............................................................................................R...
Cuchi_2          ......X......A...............-......N..S------.SD....................................................................R.E......................R...
Cuchi_3                                                           .............................................................................................R...
Cuchi_4                      .A...............-......N..........SD.............................................................................................R...
Cuchi_5_I       ..............A...............-......N..........SD......................................................................R......................R...
Cuchi_B         ..............A...............-......N..........SD.............................................................................................R...
Felix_1         .......I......A...............-......N..........SD....................................................................R.E......................R...
Felix_2                                                           ....................................................................X.X......................R...
Felix_3                                                           ...............................................................................R.............R...
Felix_4_I       ..............A...............-......N..........SD.............................................................................................R...
Felix_B         .......X......A...............-......N..........SD.............................................................................................R...
Polo_1          .......I......A...............-......N..........SD....................................................................R.E......................R...
Polo_2          .......I......A...............-......N..........SD....................................................................R.E......................R...
Polo_3                       .A....Y..........-......N..........SD....................................................................R.E......................R...
```

```
            |170                                                        |230                                                 |280
FAIDS       WDGPKMWGLRLYRTGYDPIALFTVSRQVSTITPPQAMGPNLVLPDQKPPSRQSQTGSKVATQRPQTNES--APRSVAPTTVGPKRIGTGDRLINLVQGTYLALNATDPNKTKDCWLCLVSRPPYYEGIAILGNYSNQTNPPPSCLSIPQHKL
Coco_S      ..................................R...........................S......--...........L.....................................................................
Leon_S      ....................................................F..E.............--...........L.....................................................................
Maqui_S     ...................................................L.............M...VP...P.......L.....................................................H.D..S..........
Omali_S     ...................................................P...............V.--...........L....................................................S................
Adonis_1&4  .....................................................................--S..P.......L.....................................................................
Adonis_2    .....................................................................--...........L.....................................................................
Adonis_3    .....................................................................--S..P.......L.....................................................................
Adonis_5&6  .....................................................................--...........L.....................................................................
Adonis_7    .....................................................................--...........L.....................................................................
Adonis_8    ...............................................................L.....--S..P.......L.....................................................................
Adonis_B    .....................................................................--...P.......L.....................................................................
Cuchi_1     .....................................................................--...........L.....................................................................
Cuchi_2     .....................................................................--S..P.......L.....................................................................
Cuchi_3     .....................................................................--X..X.......L.....................................................................
Cuchi_4     .....................................................................--X..X.......L..................X..................X...............................
Cuchi_5_I   .....................................................................--...........L.....................................................................
Cuchi_B     .....................................................................--...........L.....................................................................
Felix_1     .....................................................................--S..P.......L.....................................................................
Felix_2     .....................................................................--X..X.......L.....................................................................
Felix_3     .....................................................................--...........L.....................................................................
Felix_4_I   ..................................R...........................S......--...........L.....................................................................
Felix_B     .....................................................................--X..........L.....................................................................
Polo_1      .....................................................................--S..P.......L.....................................................................
Polo_2      .....................................................................--S..P.......L.....................................................................
Polo_3      .....................................................................--S..P.......L.....................................................................

            |320                                                |370                                               |420
FAIDS       TISEVSGQGLCIGTVPKTHQALCNKTQQGHTGAH--YLAAPNGTYWACNTGLTPCISMAVLNWTSDFCVLIELWPRVTYHQPEYVYTHFAKAVRFRREPISLTVALMLGGLTVGGIAAGVGTGTKALLETAQFRQLQMAMHTDIQALEESIS
Coco_S      ..................................--...................L................................................................................................
Leon_S      ..................................--....................................................................................................................
Maqui_S     ..............................A..TPL..........................L.............................G...........................................................
Omali_S     ..................................--....................................................................................................................
Adonis_1&4  ..................................--....................................................................................................................
Adonis_2    ..................................--....................................................................................................................
Adonis_3    ..................................--....................................................................................................................
Adonis_5&6  ..................................--....................................................................................................................
Adonis_7    ..................................--....................................................................................................................
Adonis_8    ..................................--....................................................................................................................
Adonis_B    ..................................--....................................................................................................................
Cuchi_1     ..................................--....................................................................................................................
Cuchi_2     ..................................--....................................................................................................................
Cuchi_3     ..................................--....................................................................................................................
Cuchi_4     ..................................--....................................................................................................................
Cuchi_5_I   ..................................--....................................................................................................................
Cuchi_B     ..................................--....................................................................................XX..............................
Felix_1     ..................................--....................................................................................................................
Felix_2     ..................................--....................................................................................................................
Felix_3     ..................................--X...................................................................................................................
Felix_4_I   ..................................--...................L................................................................................................
Felix_B     ..................................--....................................................................................................................
Polo_1      ..................................--....................................................................................................................
Polo_2      ..................................--....................................................................................................................
Polo_3      ..................................--....................................................................................................................
```

```
              |470                                                |520                                              |570
FAIDS       ALEKSLTSLSEVVLQNRRGLDILFL--QEGGLCAALKEECCFYADHTGLVRDNMAKLRERLKQRQQLFDSQQGWFEGWFNRSPWFTTLISSIMGPLLILLLILLFGPCILNRLVQFVKDRISVVQALILTQQYQQIKQYDPDRP
Coco_S      .........................--.....................................................K...............................................................
Leon_S      .........................--.....................................................K...............................................................
Maqui_S     .........................--.....................................................K...............................................................
Omali_S     .........................--.....................................................K...............................................................
Adonis_1&4  .........................--.............................................R.......K...............................................................
Adonis_2    .........................--.............................................R.......K...............................................................
Adonis_3    .........................--.............................................R.......K...............................................................
Adonis_5&6  .........................--.....................................................K...............................................................
Adonis_7    .........................--..........K.............................-----R.......K...............................................................
Adonis_8    ...................V.................................XX.................R.......K..................................................X............
Adonis_B    .........................--.....................................................K...............................................................
Cuchi_1     .........................--.............................................R.......K...............................................................
Cuchi_2     .........................--.............................................R.......K...............................................................
Cuchi_3     .........................--.............................................X.......K...............................................................
Cuchi_4     .........................--..........X.............................-----R.......K..X............................................................
Cuchi_5_I   .........................--.....................................................K...............................................................
Cuchi_B     .........................--.....X.......................................R.......K...............................................................
Felix_1     .........................--.............................................R.......K...............................................................
Felix_2     .........................--.............................................X.......K...............................................................
Felix_3     .........................--..........K.............................-----R.......K...............................................................
Felix_4_I   .........................--.....................................................K...............................................................
Felix_B     .........................--..........X..........................................K...............................................................
Polo_1      .........................--.............................................R.......K...............................................................
Polo_2      .........................--.............................................R.......K...............................................................
Polo_3      .........................--.............................................R.......K...............................................................
```
